# Sequence, Structure and Functional space of *Drosophila de novo* proteins

**DOI:** 10.1101/2024.01.30.577933

**Authors:** Lasse Middendorf, Bharat Ravi Iyengar, Lars A. Eicholt

**Affiliations:** Institute for Evolution and Biodiversity, University of Muenster, Muenster, Germany

**Author notes:** corresponding authors: Lars A. Eicholt & Bharat Ravi Iyengar, Huefferstrasse 1, 48149 Muenster, Germany, &. **Declaration of Interests** The authors declare no competing interests.

**Keywords:** *de novo* proteins, protein function, structural comparison, protein structure, structure predictions, sequence space

## Abstract

During *de novo* emergence, new protein coding genes emerge from previously non-genic sequences. The *de novo* proteins they encode are dissimilar in composition and predicted biochemical properties to conserved proteins. However, many functional *de novo* proteins indeed exist. Both identification of functional *de novo* proteins and their structural characterisation are experimentally laborious. To identify functional and structured *de novo* proteins *in silico*, we applied recently developed machine learning based tools and refined the results for *de novo* proteins. We found that most *de novo* proteins are indeed different from conserved proteins both in their structure and sequence. However, some *de novo* proteins are predicted to adopt known protein folds, participate in cellular reactions, and to form biomolecular condensates. Apart from broadening our understanding of *de novo* protein evolution, our study also provides a large set of testable hypotheses for focused experimental studies on structure and function of *de novo* proteins in *Drosophila*.

## Introduction

Once considered impossible [Zuckerkandl, 1975, Jacob, 1977], many lines of evidence suggest that functional proteins can emerge from random sequences that have not been subjected to several generations of evolution [Keefe and Szostak, 2001, Hecht et al., 2004, Babina et al., 2023]. For example, high throughput selection experiments with a large number of random sequences have shown, that some random proteins can mitigate auxotrophy [the inability to metabolize nutrients; Knopp et al., 2021], provide resistance against toxins [Frumkin and Laub, 2023], and even catalyze biochemical reactions [Chao et al., 2013, Yamauchi et al., 2002]. In accordance with the fact that protein folding is often a critical requirement for protein function, many random proteins have been also shown to have secondary structures [Davidson and Sauer, 1994, Davidson et al., 1995, Tretyachenko et al., 2017, Surdo et al., 2004, Mansy et al., 2007]. *De novo* emergence is a phenomenon through which novel protein coding genes arise from non-genic regions of the genome [Tautz and Domazet-Lošo, 2011, Carvunis et al., 2012, Oss and Carvunis, 2019, Vakirlis et al., 2020a, Bornberg-Bauer et al., 2021, Schmitz and Bornberg-Bauer, 2017]. The *de novo* proteins thus emerged have been considered to be the natural equivalent of random sequences, because they emerge from supposedly “random” intergenic regions, and some of their predicted properties such as length, structural disorder and aggregation propensity, resemble that of random proteins, more than that of conserved proteins [Heames et al., 2023, Bornberg-Bauer et al., 2021, Ángyán et al., 2012, Bhave and Tautz, 2021, Castro and Tautz, 2021, Middendorf and Eicholt, 2024, Aubel et al., 2024]. For example, *de novo* proteins in *Drosophila*, are predicted to be more disordered than conserved proteins [Heames et al., 2020, Middendorf and Eicholt, 2024, Peng and Zhao, 2023], which can be partially explained due to higher GC content of the former [Landry et al., 2015, Zheng and Zhao, 2022]. While the structure of large sets of *de novo* proteins have been computationally analyzed [Schmitz et al., 2018, Heames et al., 2020, Peng and Zhao, 2023, Basile et al., 2017, Chen et al., 2023, Vakirlis et al., 2020b], the structures of only four *de novo* proteins have been experimentally approximated [Lange et al., 2021, Bungard et al., 2017, Baalsrud et al., 2018, Matsuo et al., 2021]. Determining the function of *de novo* genes and proteins is another challenging task. It involves identifying the cell types and stages in which *de novo* proteins may be involved and testing their phenotypic effects using genetic tools [Chen et al., 2010a, Gubala et al., 2017, Lange et al., 2021, Reinhardt et al., 2013]. Nonetheless, functional *de novo* proteins indeed exist and have been identified in organisms as diverse as insects, plants (*Arabidopsis thaliana*), fungi (*Saccharomyces cerevisae*), arctic codfish, mice (*Mus musculus*) and humans (*Homo sapiens*) [McLysaght and Guerzoni, 2015, Li et al., 2009, Cai et al., 2008, Chen et al., 2010a, Gubala et al., 2017, Lange et al., 2021, Zhuang et al., 2019, Reinhardt et al., 2013, Heinen et al., 2009, Li et al., 2010a, Xie et al., 2019, Li et al., 2014, Vakirlis et al., 2022, Linnenbrink et al., 2024, Klasberg et al., 2018, Li et al., 2010b, Matsuo et al., 2021, Rivard et al., 2021, Begun et al., 2007]. Experimental structure determination is a laborious process that cannot be performed in a high throughput manner. This is especially difficult for *de novo* proteins because of high aggregation propensity and low solubility *in vitro* [Eicholt et al., 2022]. Despite the increasing numbers of solved structures, novel structures, whether they be folds or domains, were rarely ever found [Grant et al., 2004, Levitt, 2009, Tóth-Petróczy and Tawfik, 2014]. However, the recent advancements in highthroughput structure predictions through computational techniques, have led to discovery of novel folds [Durairaj et al., 2023]. Since *de novo* proteins are void of ancestry from conserved protein families, they could provide rare structural novelty [Bornberg-Bauer et al., 2021]. From another perspective, the occurrence of conserved or ancient structural folds in *de novo* proteins could suggest a high level of evolutionary accessibility in sequence space. This might explain the emergence of these folds during the early stages of protein evolution [Lupas et al., 2001, Kopec and Lupas, 2013, Alva et al., 2010, 2015, Romero Romero et al., 2016]. A protein’s structure can provide some clues about its function [Orengo et al., 1999]. For example, one can reasonably guess the function of an uncharacterized protein by comparing its structure to that of a known functional protein [Nomburg et al., 2024]. Although, protein function is often attributed to its structure, and unfolded proteins were assumed to be toxic, many studies show that disordered proteins can be functional [Deiana et al., 2019, Jemth et al., 2018, Ali and Ivarsson, 2018]. For example, disordered proteins can help form intracellular condensates (or membrane less organelles) that have been shown to play a major role in the cellular physiology of diverse organisms [Lin et al., 2017, Hyman et al., 2014]. Because *de novo* proteins could be a source of novelty, with regards to both structure and function, we aimed to understand their structures and possible functions through computational analyses. To this end, we studied a previously characterized set of 2510 putative *de novo* proteins from the *Drosophila* clade [Heames et al., 2020, Middendorf and Eicholt, 2024]. We used a multi-faceted approach analyze these *de novo* proteins. First, we used Foldseek [van Kempen et al., 2023] to find experimentally known protein structures [Protein Data Bank, Berman et al., 2000] and predicted protein structures [AlphaFold database, Varadi et al., 2021] that are similar to the AlphaFold2 (AF2) [Jumper et al., 2021] predicted structures of our *de novo* proteins. Second, we predicted the functions of our *de novo* proteins using DeepFRI [Gligorijević et al., 2021], a machine learning-based tool that predicts functional annotations (gene ontology terms) using protein structure and sequence features. Because many of our *de novo* proteins were predicted to be disordered *de novo* proteins, we hypothesized that they could form biomolecular condensates [Uversky, 2017]. To test this hypothesis, we predicted the condensate forming propensity of our *de novo* using PICNIC [Hadarovich et al., 2023], an algorithm that is based on predicted structure (AlphaFold2), predicted disorder (IUPred2A), as well as sequence complexity. Understanding the condensate forming behavior of *de novo* proteins would elucidate their potential involvement in the formation of membraneless organelles, offering an evolutionarily and biophysically feasible mechanism for their integration with the cellular physiology. Finally, we mapped the *de novo* proteins on the protein sequence space in relation to random and conserved proteins. To this end, we used protein language models that can predict several biophysical features from sequences, embedding their abstracted properties in the form of numerical values [Lin et al., 2023]. Our method allowed us to map different sequences with better resolution than by the analyses of individual properties separately [Weidmann et al., 2019, Agozzino and Dill, 2018, Heames et al., 2023, Aubel et al., 2024]. With these multi-faceted analyses we found that some *de novo* proteins can indeed adopt structures similar to known proteins and can have possible cellular activities including localization to specific organelles. We also found that some *de novo* proteins are likely to form biomolecular condensates. However, with our language model analysis we found that the majority of *de novo* proteins look distinct from conserved proteins of similar length, and resemble more the random proteins. Overall, our work enhances our understanding of how *de novo* proteins can not only develop features already known to the living systems, but can also be a source for evolutionary novelty.

## Results

### A few *de novo* proteins can indeed adopt known structures

To understand if *de novo* proteins can form known protein structures, we compared their predicted structure to that of conserved proteins. Recent studies have shown that structure predictions are not very reliable for *de novo* proteins [Middendorf and Eicholt, 2024, Aubel et al., 2023, Liu et al., 2023], and that many predicted structures are also thermodynamically unstable [Peng and Zhao, 2023]. Therefore, we refined the predicted structures of *Drosophila de novo* proteins from our previous study Middendorf and Eicholt [2024], using molecular dynamics simulations, performing 3 replicate simulations per protein for 100ns. We thus refined the predicted structures of 1,468 *de novo* proteins. Our MD simulations suggest that most *de novo* proteins exhibit structural flexibility, as indicated by the large root mean square deviation (RMSD) values (Figure Figure 1A and Figure S3). Next, we searched for conserved proteins that have predicted structures similar to those of *de novo* proteins, using Foldseek [van Kempen et al., 2023]. Specifically, with MD refined structures as queries, and the AFDB50 [Varadi et al., 2021] as the target, we observed that the majority of *de novo* proteins did not have a significant structural similarity to the conserved proteins in AFDB50 (TM score <0.5, Figure 1B). This was also the case for AF2 predicted structures of *de novo* and random proteins without MD simulations (Figure S1 and Figure S2). This observation, supports the *de novo* status of our proteins, aligning with the notion that structure is more conserved than sequence [Illergård et al., 2009]. To investigate whether these *de novo* proteins can adopt known structures, we performed structural mapping of *de novo* proteins with experimentally validated structures in the Protein Data Bank (PDB) [Berman et al., 2000], using Foldseek. We then extracted the ECOD domain annotations for matches found in the PDB [Cheng et al., 2014]. Out of the 1,468 *de novo* proteins analyzed, 42 showed structural alignment with proteins having an architecture annotation in ECOD (Figure 1C). Prior to MD simulation, 119 predicted structures were mappable to PDB structures (Figure S1). Figure 1D presents examples of these findings consisting of a structurally unalignable *de novo* protein, one similar to an SH3 fold, and another resembling an HTH fold. Both SH3 and HTH folds are considered highly conserved and ancient folds [Kishan and Agrawal, 2005, Alvarez-Carreño et al., 2021, Rosinski and Atchley, 1999, Grishin, 2000]. These three example proteins have emerged less than 5 million years ago (mya) [Heames et al., 2020]. Overall, our structure search analysis shows that, while most *de novo* proteins are likely to have novel or uncommon structures, a minority of them can indeed adopt well known protein structures.

**Figure 1:**
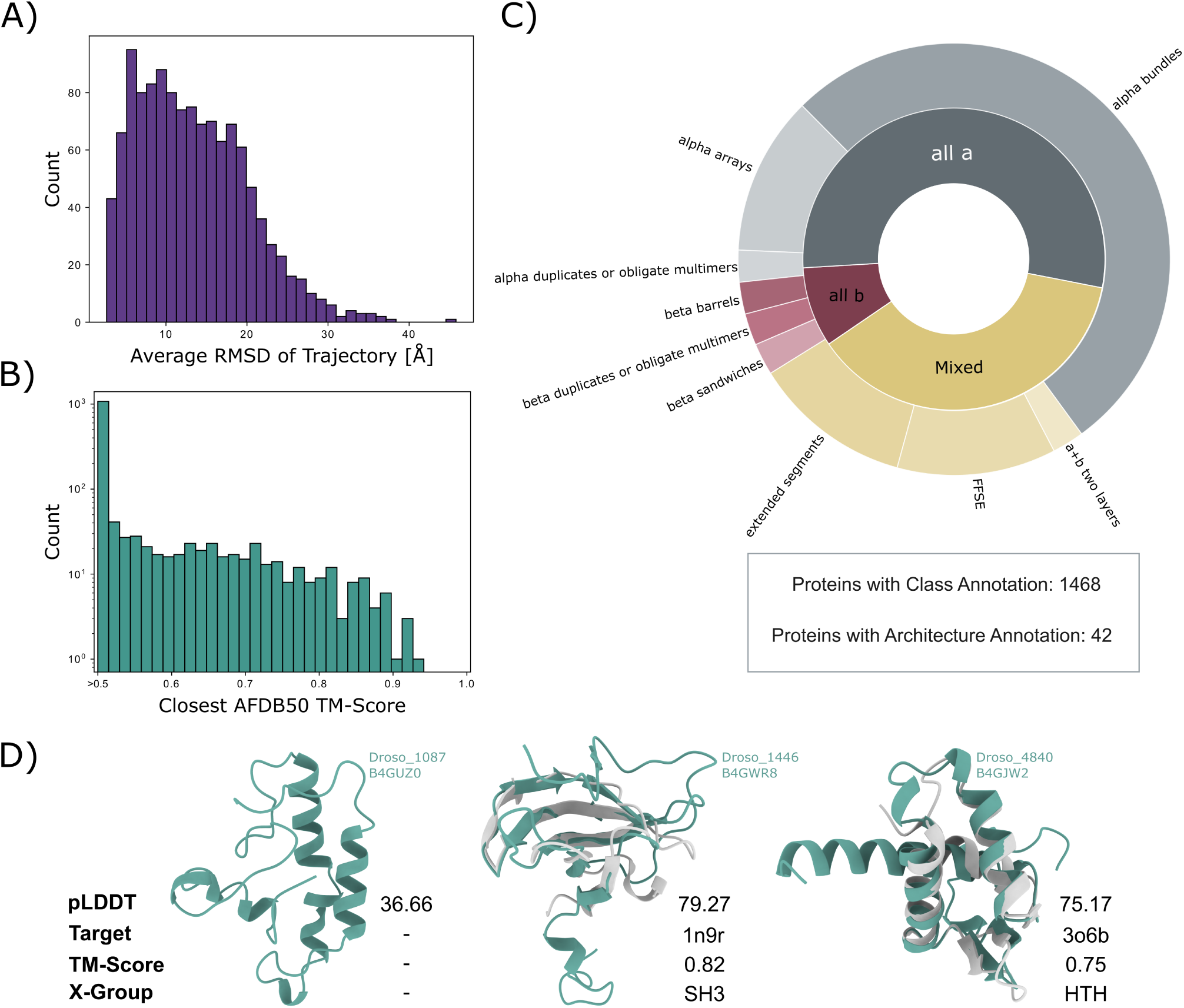
Structural diversity of *de novo* evolved proteins. (**A**) Distribution of the average root mean square deviation (RMSD, horizontal axis) per MD simulation trajectory. We display the average RMSD of three MD simulation replicates per *de novo* protein, only for proteins with i) less than 30% disorder predicted by flDPnn, and ii) less than 95% of their residues annotated as α-helices via DSSP (1468 of 2510 proteins). (**B**) Distribution of the TM-score (horizontal axis) for the mapping of *de novo* proteins (MD-refined structures) to the most similar protein structure in the AlphaFold database (AFDB50), excluding proteins from *Drosophila*. TM-scores below 0.5 indicate no similarity to any protein structure in the AlphaFold database. (**C**) Structural classification of *de novo* proteins. We assigned a structural class to each of the 1468 *de novo* proteins based on the DSSP annotations of their predicted structures (inner circle). To identify annotated protein domains in *de novo* proteins, we aligned their MD refined structures to structures in the PDB. We assigned each *de novo* protein with the ECOD domain of its highest scoring hit from the PDB, given the TM-score was greater than 0.5 and the alignment covered at least 80% of the PDB target. We assigned the 42 *de novo* proteins, that qualified the above criteria, with an ECOD domain from multiple domain architectures (outer circle). (**C**) Examples of *de novo* proteins without structural similarity to proteins in the AlphaFold database (Droso_1087), or with similar structure to an ECOD X-group (Droso_1446 & Droso_4840; aligned with their closest hit in the PDB).

### Some *de novo* proteins may bind to nucleic acids, and are predicted to have enzymatic activities

Information on biological activities and functions, is available for only a handful of *de novo* proteins [Bornberg-Bauer et al., 2021, Weisman, 2022]. The existence and gain of biological activity would be critical factor determining the evolutionary fixation of *de novo* proteins. However, the lack of homology, makes functional annotation challenging. Therefore, we used DeepFRI to functionally annotate *de novo* proteins with Gene Ontology (GO) terms. Unlike homology based techniques, DeepFRI combines a protein language model, trained on the sequences of PFAM domains, and a graph convolutional network that represents amino acid interactions derived from protein structure [Gligorijević et al., 2021]. DeepFRI is also trained on the GO terms associated with different structures. We did not filter protein sequences according to any structural criteria, because DeepFRI can de-noise predicted protein structures [Gligorijević et al., 2021]. We summarized and clustered the predicted GO terms based on their semantic similarity, and projected them in a 2-dimensional semantic space using REVIGO [Supek et al., 2011] (Figure 2A & B). We identified these GO term clusters visually and manually annotated them based on the GO terms within the cluster. We performed this analysis for both *de novo* and random proteins. With our analysis, we found that a small fraction of *de novo* and random proteins could be confidently annotated with GO terms for all the three GO classes (Molecular Function, Biological Process, and Cellular Component; Figure 2C). The GO term class *Cellular Component* had the highest fraction of confident predictions with *≈*31% and *≈*17% for *de novo* and random proteins, respectively. However, we could not find any overarching GO terms within the cellular component category, for both *de novo* and random proteins. This suggests that both these kind of proteins can localize to many different cellular compartments. Specifically, we found that these proteins, can possibly localize to the following compartments: nucleus (GO:0005634), mitochondrion (GO:0005739), vesicles (GO:0031982), and membranes (GO:0016020).

**Figure 2:**
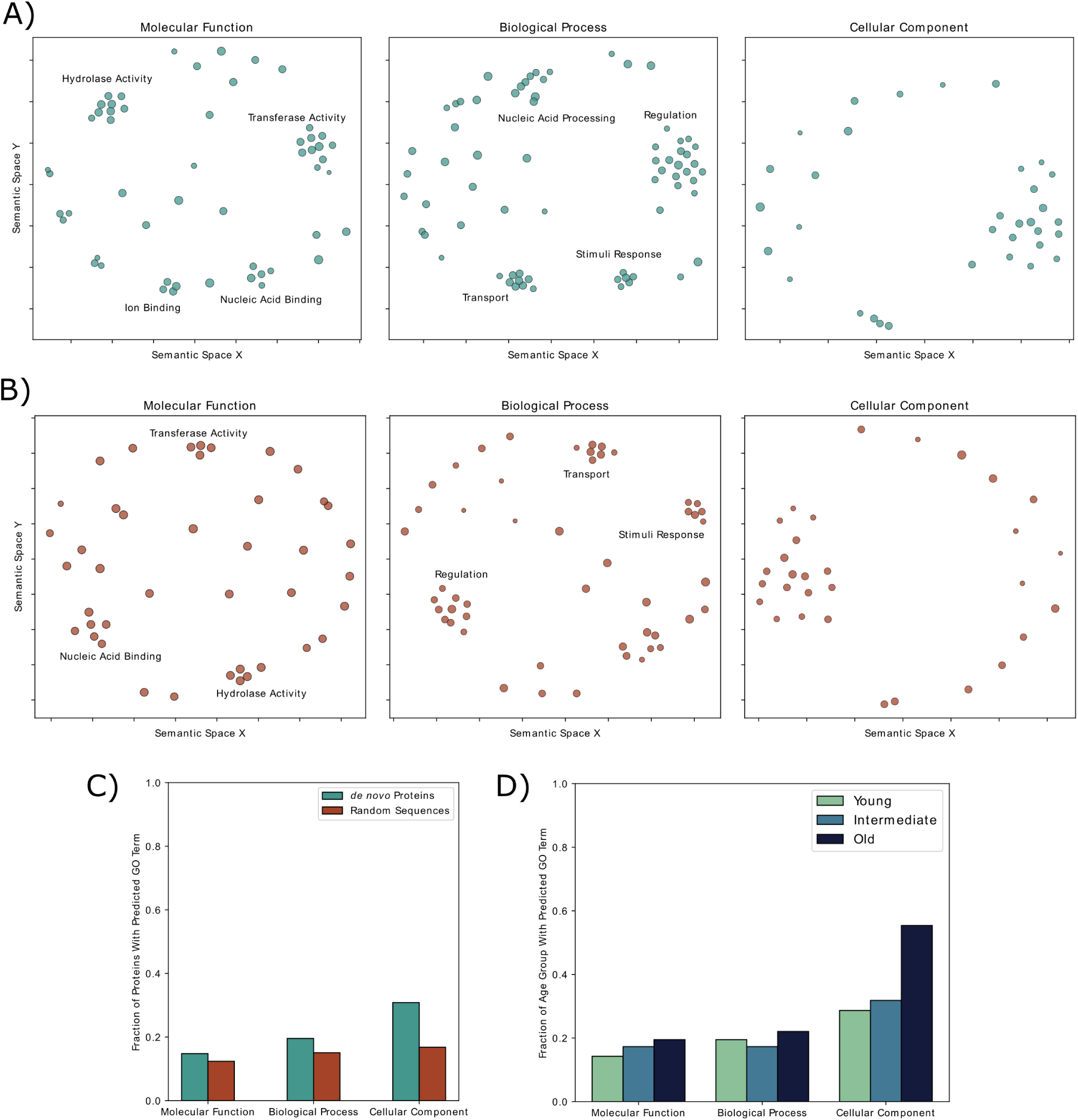
GO terms of random and *de novo* proteins predicted with DeepFRI. We predicted GO terms of *de novo* proteins (**A**) and random sequences (**B**) with DeepFRI and clustered them based on semantic similarity with REVIGO. We visually identified GO term clusters manually annotated with a generic term that describes all the GO terms within the respective cluster. (**C**) Fraction of *de novo* and random proteins (vertical axis) predicted with a GO term per GO term category (horizontal axis). (**D**) Fraction of *de novo* proteins in different age groups (vertical axis) with a predicted GO term (horizontal axis). *Old de novo* proteins were significantly more often annotated with a GO term in the *Cellular Component* category than expected by chance (Pearson’s *χ*^2^-Test; *P <* 10*^−^*^10^).

Both *de novo* proteins and random sequences both show a broad variety of GO terms in other two GO classes with only a few prominent clusters within the semantic space (Figure 2A & B). Interestingly, *de novo* proteins and random sequences appear to have similar molecular functions and to be involved in similar categories of biological processes. Regarding their molecular function, they both showed multiple GO terms in relation to “hydrolase activity”, “transferase activity”, and “nucleic acid binding”. The biological processes in which *de novo* proteins and random sequences are both predicted to be involved were “stimuli response”, “regulation” and “transport”. Next, we analyzed the impact of evolutionary age on functional annotation using GO terms. As young *de novo* proteins were more frequent than older proteins in the dataset, we normalized the number of proteins with predicted GO terms to the number of proteins in the respective age group. In all three categories of GO terms, the oldest *de novo* proteins (emerged >30 Mya) were more often predicted with a GO term, than younger proteins (Figure 2D). Only for the GO term category *Cellular Component*, old *de novo* proteins were annotated more frequently than expected by chance (Pearson’s *χ*^2^-Test; *P <* 10*^−^*^10^).

### Subset of *de novo* proteins may form biomolecular condensates

Biomolecular condensates are membraneless compartments formed by proteins via liquid-liquid phase separation, and are involved in several biological processes such as stress response and regulation of transcription [Tsang et al., 2020, Hyman et al., 2014]. We observed that that GO terms concerning RNA binding, transferase activity, and hydrolase activity that predicted for *de novo* proteins (Figure 2), are also important features of condensate-forming proteins [Hadarovich et al., 2023]. Therefore, we predicted the propensity of *de novo* proteins for condensate-formation. To this end, we used another prediction tool called PICNIC [Hadarovich et al., 2023]. However, PICNIC uses AF2 predicted structures and a disorder prediction tool IUPred2A, to predict condensate formation propensity. It has been shown, that both AF2 and IUPred can make qualitatively discordant predictions of *de novo* proteins [Middendorf and Eicholt, 2024, Aubel et al., 2023]. Therefore, we performed additional analyses to ensure a high-confidence prediction of condensate-forming *de novo* proteins (Figure 3A). Specifically, we retrieved 175 known condensate-forming conserved proteins from the CD-CODE database [Rostam et al., 2023] and used them as a positive control dataset. For all these proteins, we calculated the sequence features that are associated with the biological function of their intrinsically disordered regions, e.g. amino acid homorepeats, sequence complexity, and net charge [Zarin et al., 2021]. We clustered sequences based on these sequence features using Uniform Manifold Approximation and Projection (UMAP) [McInnes et al., 2018], a commonly non-linear dimensionality reduction tool (in contrast to principal component analysis, which is linear; Figure 3B). We identified seven clusters of different sizes. Of these, cluster 1 and cluster 3 contained most proteins (88.6%) of the CD-CODE database that we used in our analysis (Figure 3C). The *de novo* proteins in cluster 1 and cluster 3 with a PICNIC score greater than 0.5 can be considered high-confidence condensate forming proteins, because they are not only predicted by PICNIC according to its own criteria, but they also have a similar sequence composition as experimentally validated condensate-forming proteins. In total, we identified 63 such high-confidence condensate-forming *de novo* proteins. We next analysed the age groups of these condensate forming *de novo* proteins. When normalized by the number of proteins per age group, we found intermediate and old *de novo* proteins to be 5.9– and 6.6-fold more often predicted to form condensates than young *de novo* proteins, respectively (Figure 3D). Furthermore, intermediate and old *de novo* proteins contained significantly more high-confidence condensate-forming proteins than expected by chance (Pearson’s *χ*^2^-Test; *P <* 5 *×* 10*^−^*^54^).

**Figure 3:**
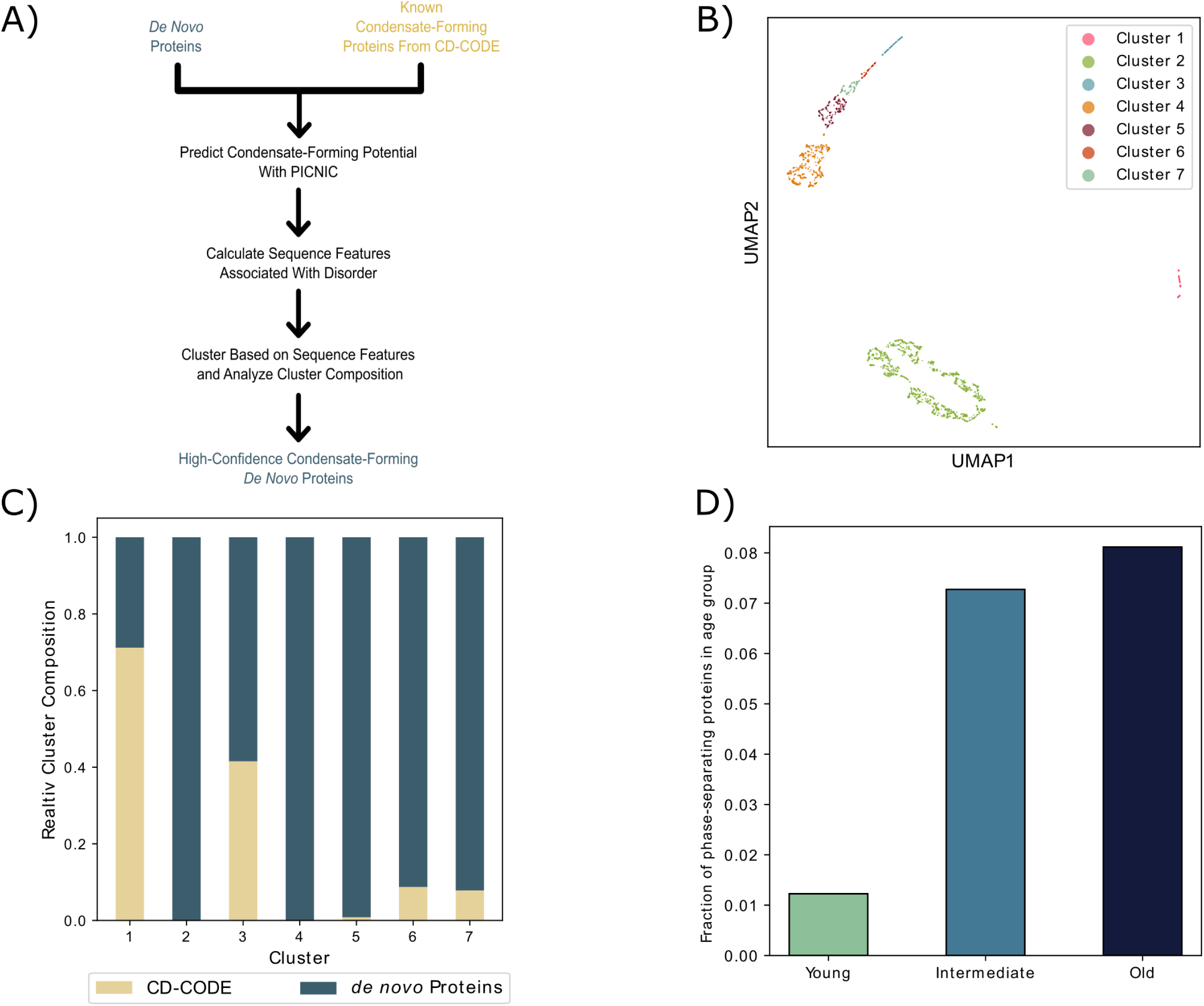
Identification of condensate-forming *de novo* proteins. (**A**) Workflow for the identification of condensate-forming *de novo* proteins. We predicted condensate-forming potential of *de novo* proteins and known condensate-forming proteins from the CD-CODE database with PICNIC. For both groups of proteins, we calculated the sequence features associated with the functions of intrinsically disordered regions were calculated. Subsequently, we clustered all proteins based on these sequence features using hdbscan, and the analyzed the clusters for their constituent proteins. (**B**) Clusters of *de novo* proteins and known condensate-forming proteins based on sequence features associated with the function of intrinsically disordered proteins. (**C**) Constitution of the identified clusters based on protein type. We classified the 63 *de novo* proteins from clusters 1 and 3 were as high-confidence condensate-forming proteins. (**D**) Fraction of *de novo* proteins from the respective age groups that were classified as high-confidence condensate-forming proteins. The age groups *Intermediate* and *Old* contained significantly more high-confidence condensate-forming proteins than expected by chance (Pearson’s *χ*^2^-Test; *P <* 5 *×* 10*^−^*^54^).

### Protein language models show that *de novo* and conserved proteins occupy distinct regions of the sequence space

Although we found that some *de novo* proteins may be structurally similar to known proteins, we don’t yet know if evolutionary origin indeed determines the structural properties of a protein. Indeed, many studies have compared a handful of features such as structural disorder, protein composition, and aggregation propensity between *de novo* and conserved proteins [Knowles and McLysaght, 2009, Ekman and Elofsson, 2010, Landry et al., 2015, Wilson et al., 2017, Vakirlis et al., 2018, Klasberg et al., 2018, Schmitz et al., 2018, Heames et al., 2020, 2023, Peng and Zhao, 2023, Middendorf and Eicholt, 2024]. However, these analyses may not provide reliable inferences because they use tools depending on limited data (e.g. TANGO/IUPred) [Fernandez-Escamilla et al., 2004, Erdős et al., 2021], and because the different features are analysed in isolation. Language models use machine learning to analyse several hidden parameters (and their interactions) simultaneously using sequence information alone. Indeed, protein language models have proved extremely adept at predicting and designing protein structures [Heinzinger et al., 2019, Madani et al., 2023, Alley et al., 2019, Chowdhury et al., 2022, Ferruz and Höcker, 2022, Ferruz et al., 2023, Lin et al., 2023].Therefore, we used the ESM2 protein language model to compare the three different kinds of protein sequences in our dataset (random, *de novo* and conserved proteins). Specifically, we generated a numerical vector for each protein sequence using the ESM2 language model with 650 million parameters (ESM2-650M) [Lin et al., 2023]. Each vector contains 1280 elements, that denote an abstraction of different sequence features predicted by the model. We used UMAP [McInnes et al., 2018] to visualize the protein sequences in sequence space, and found that *de novo*, random, and conserved proteins indeed occupy distinct regions in the sequence space (Figure 4). To quantify these observations, we calculated the Manhattan distance (or L1 norm) between every pair of protein numerical sequences, a method particularly effective for multidimensional data with potential extreme outliers [Barrodale, 1968]. Our findings indicate that the distances between *de novo* and conserved proteins are generally larger than those between sequences within each of these categories (one-sided Mann-Whitney U test; *P <* 10*^−^*^99^). We also found that the distances between the *de novo* and conserved proteins are generally larger than the distances between the *de novo* and the random proteins (one-sided Mann-Whitney U test; *P <* 10*^−^*^99^). The generated random proteins were based on the same length and amino acid distributions as the *de novo* proteins [Middendorf and Eicholt, 2024, Heames et al., 2023]. Therefore, the nearness between these two sets of protein sequences could be an artifact of our method. To verify if this is the case, we generated random protein sequences with same distribution of composition as our conserved sequences. We found that *de novo* proteins were closer to these new random proteins than with conserved proteins (one-sided Mann-Whitney U test; *P <* 10*^−^*^99^; Figure S4). Overall our analyses suggest that despite certain structural similarities, *de novo* proteins are, distinct from conserved proteins at the sequence level, and bear a closer resemblance to random sequences.

## Discussion

Most proteins can be grouped into families based on their sequence similarity, evolutionary ancestry, structural folds, and biochemical functions [Chothia, 1992]. *De novo* proteins are exceptions as they do not belong to any established protein family, because they not only originate from nongenic DNA sequences (lack of ancestry), but also lack sequence and structural homology to other proteins [Bornberg-Bauer et al., 2021, Schlötterer, 2015]. This makes it challenging to annotate functions to *de novo* proteins based on our knowledge of conserved proteins. Despite their dissimilarity with known proteins, *de novo* proteins have been shown to perform biological functions and improve the survival and fitness of the organisms that express them [McLysaght and Guerzoni, 2015, Li et al., 2009, Cai et al., 2008, Chen et al., 2010a, Gubala et al., 2017, Lange et al., 2021, Zhuang et al., 2019, Reinhardt et al., 2013, Heinen et al., 2009, Li et al., 2010a, Xie et al., 2019, Li et al., 2014, Chen et al., 2010b]. Advanced computational methods using deep learning have been able to solve problems at an unprecedented scale. For example, AlphaFold2 resulted in an exponential increase in the number of computationally predicted protein structures [Varadi et al., 2021] Therefore, we applied some of these deep learning based tools to elucidate the possible structure and function of *de novo* proteins.

First, we searched for conserved proteins that may be structurally similar to *de novo* proteins using Foldseek. Most *de novo* proteins did not bear a significant resemblance to known protein structures, in accordance with their non-genic evolutionary origin, and distinctiveness of their sequence and biophysical properties as shown by previous studies [Heames et al., 2023, Aubel et al., 2024]. This lack of resemblance could exist because *de novo* proteins are highly disordered [Middendorf and Eicholt, 2024, Peng and Zhao, 2023], and can contain rare secondary structures like 3_10_– or *π*-helices [Chen et al., 2023], that could make structural alignment complicated.

While we attempted to refine AF2 predicted structures of *de novo* proteins through molecular dynamics (MD) simulations, it is important to note that many *de novo* proteins may reside in nonaqueous environments such as cell membranes (Figure Figure 2) [Vakirlis et al., 2020b], may only fold upon interaction with other proteins [Chen et al., 2023], and may be part of multimers [Lynch, 2012, Schulz et al., 2022, Jayaraman et al., 2022, Malik et al., 2024]. We did not consider all these possibilities in our MD simulations due to computational limitations. Nonetheless, the majority of individual *de novo* proteins were predicted to be disordered or, if structured, to predominantly form simple α-helices [Heames et al., 2023, Middendorf and Eicholt, 2024, Aubel et al., 2024, Peng and Zhao, 2023], a trend attributed to many *de novo* proteins being too short to form globular structures [Aubel et al., 2024, Shen et al., 2005]. Our current study corroborates these observations. The frequent emergence of single α-helices in *de novo* proteins can be attributed to the lower stereo-chemical and thermodynamical requirements of α-helices [Barlow and Thornton, 1988, Greenwald and Riek, 2012]. On rare occasions where *de novo* proteins exhibit structural configurations beyond single α-helices, they can resemble common and ancient folds such as SH3 or HTH (Figure 1D). This observation implies that these widespread evolutionary folds, which are evolutionary successful and easily tolerated by cells, are more accessible in sequence space [Taverna and Goldstein, 2000, Shakhnovich et al., 2005, Goldstein, 2008], even for sequences that have not been shaped by millions of generations of evolution. Despite identifying some *de novo* proteins with structural homology to existing structures, we did not find any novel folds among our candidate proteins, unlike other studies that investigated a much larger set of proteins [Durairaj et al., 2023] (Figure 1B & D).

By employing the deep learning based functional annotation tool, DeepFRI [Gligorijević et al., 2021], we found that *de novo* proteins are associated with a wide array of Gene Ontology (GO) terms, spanning all three GO categories, with several distinct clusters emerging within the semantic field. We show that *de novo* proteins, despite their recent emergence and lack of evolutionary ancestry, are more often predicted to be functional than a comparable random set of sequences (Figure 2C). Their involvement in a range of molecular functions (like hydrolase activity, transferase activity, and nucleic acid binding) and biological processes (such as stimuli response, regulation, and transport) underscores their potential impact on the cellular physiology. Interestingly, the similarity in molecular functions and involvement in biological processes between *de novo* proteins and random sequences could imply a certain level of functional redundancy in the sequence space. This observation might suggest that the emergence of function from novel proteins, even through random sequences, could be a more probable phenomena than previously thought. Finally we emphasize that, while efforts to deduce protein function based on structural similarity is ongoing [Nomburg et al., 2024, Gligorijević et al., 2021], numerous instances exist where proteins with similar structures perform different functions, and *vice versa* [Finkelstein et al., 1993, Govindarajan and Goldstein, 1996, Galperin et al., 1998, Martin et al., 1998].

The association of *de novo* proteins with biophysical reactions such as RNA binding, and biochemical reactions similar to transferases, and hydrolases, presents an intriguing avenue for understanding their functional capacities and evolutionary significance. This is especially interesting because RNA binding and hydrolase-activity are thought to be conferred even by primordial folds [Seal et al., 2022, Weil-Ktorza et al., 2023, Vyas et al., 2021, Longo et al., 2022], and could possibly been important during origin of life. Both these molecular activities, and a highly disordered structure, are also exhibited by condensate-forming proteins [Hadarovich et al., 2023]. Therefore, we investigated the possibility of *de novo* proteins to be involved in formation of biomolecular condensates. Biomolecular condensates, formed through liquid-liquid phase separation by proteins, are critical in various biological processes and such a propensity exists even for proteins with ancient and simple folds [Longo et al., 2020]. The use of PICNIC [Hadarovich et al., 2023] to predict the involvement of *de novo* proteins in biomolecular condensates represents an innovative approach, albeit with limitations. The reliance on AlphaFold2 predictions and IUPred2A as input requirements, introduces a degree of uncertainty, especially given the discordant predictions of these tools between *de novo* and conserved proteins [Middendorf and Eicholt, 2024]. This necessitated further analysis to establish a high-confidence set of condensate-forming *de novo* proteins, leveraging the CD-CODE database [Rostam et al., 2023] as a reference.

The identification of clusters based on sequence features associated with intrinsically disordered regions of proteins is particularly noteworthy. The fact that clusters 1 and 3, which have a high fraction of members from the CD-CODE database, include *≈*12% of all *de novo* proteins with a PICNIC score greater than 0.5, is compelling. It suggests that these *de novo* proteins not only have the potential to form condensates but also share sequence composition with experimentally validated condensate-forming proteins. The discovery of 63 high-confidence condensate-forming *de novo* proteins contributes to our understanding of the functional diversity of these proteins. This finding expands the realm of *de novo* protein functionality beyond traditional views, indicating their potential involvement in complex cellular mechanisms like phase separation. Considering that phase separation is involved in spermatogenesis [Kang et al., 2022, Parvinen, 2005], and that *de novo* proteins show biased expression in testis [Levine et al., 2006, Heames et al., 2020, Zhao et al., 2014, Palmieri et al., 2014, Peng and Zhao, 2023, Nyberg and Carthew, 2017, Kondo et al., 2017, Neme and Tautz, 2013], being involved in biomolecular condensates suggests a possible mechanism by which *de novo* proteins could play a role in spermatogenesis [Lange et al., 2021, Gubala et al., 2017, Rivard et al., 2021]. Moreover, our analysis of the age groups of these *de novo* proteins revealed that intermediate and old *de novo* proteins are significantly more likely to form condensates than their younger counterparts. This observation is intriguing as it could imply two scenarios. First, as *de novo* protein evolve and mature, they acquire and refine their ability to participate in cellular processes like biomolecular condensation and thereby their function. Under this scenario, the *de novo* proteins could be positively selected. Second, the ability to form biomolecular condensates could minimize toxic protein aggregation, and could protect *de novo* proteins from being purged by negative selection.

To understand if *de novo* proteins can indeed be a source of evolutionary novelty, we analyzed their distribution in the protein sequence space relative to that of conserved and random proteins, using the protein language model ESM2-650M. Our analysis shows that *de novo* proteins, arisen from non-coding sequences, have unique sequence characteristics that distinguish them from conserved proteins, but more similar to random proteins, as hypothesized before [Bornberg-Bauer et al., 2021]. Nevertheless, some *de novo* proteins indeed had a conserved protein, closely located to them in the sequence space (Figure 4). Together with our Foldseek analysis, this observation indicates an inherent capacity of amino acid sequences to adopt structures, and that a broad spectrum of sequence space is capable of evolving into foldable proteins [Tretyachenko et al., 2017, Heames et al., 2023, Aubel et al., 2024].

**Figure 4:**
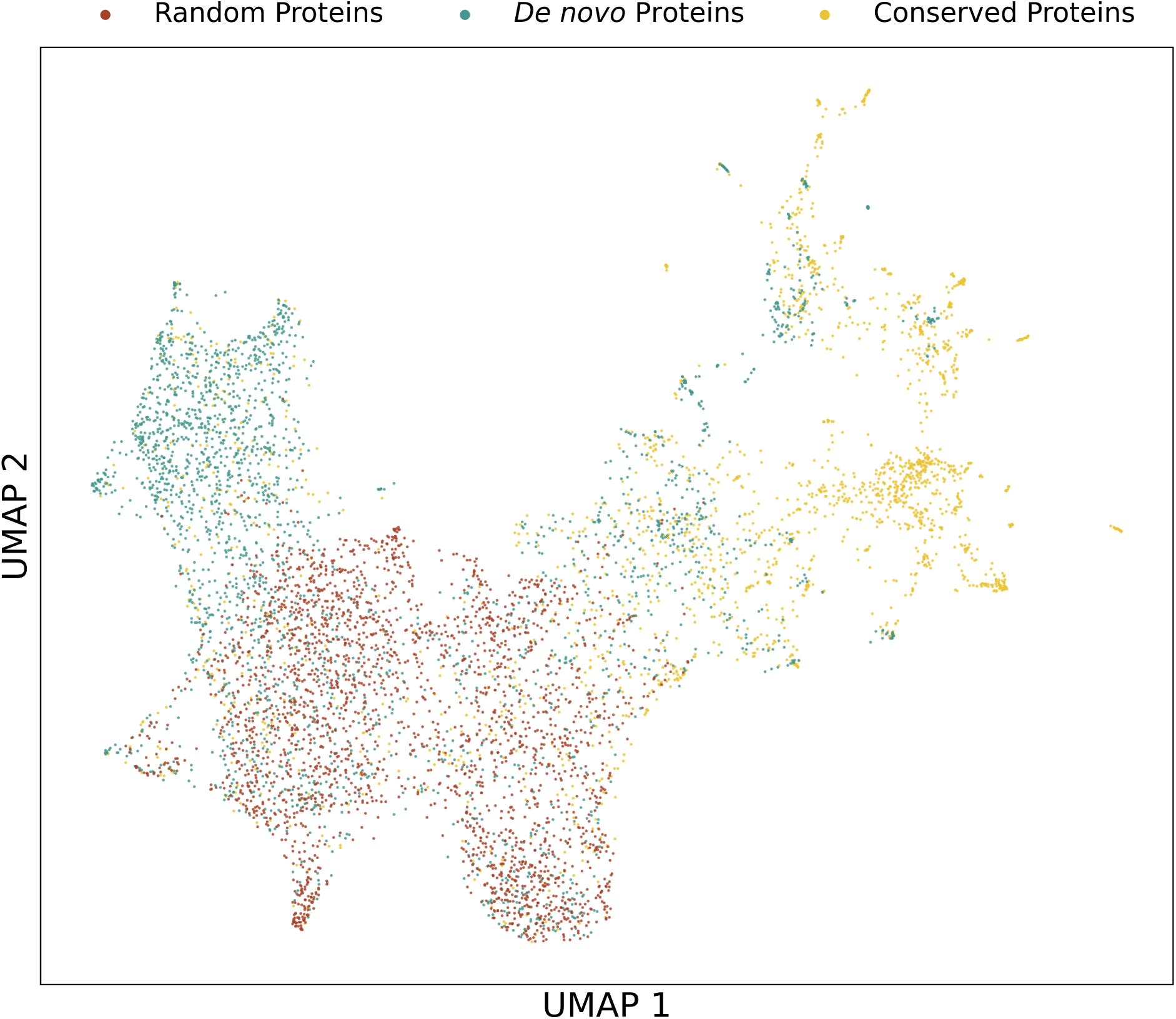
Location of our protein sequences in the sequence space. We used the protein language model ESM2-650M to generate a numerical representation of the *de novo*, random and conserved proteins sequences. We projected and plotted these numerical sequences into a two dimensional space using UMAP.

Our analysis is based on computational tools, which are always prone to some level of erroneous predictions. Furthermore, many of the deep learning based tools have not been trained on *de novo* proteins and can possibly make biased predictions [Middendorf and Eicholt, 2024]. Therefore, our study may not provide exact and perfect answers to the different open questions about *de novo* proteins. All computational predictions need experimental validation. Experimental studies, especially on *de novo* proteins are bottlenecked by serendipity, and labor intensive techniques that are not fully optimized for proteins with such an unusual biochemistry [Eicholt et al., 2022]. However, our exhaustive approach can help guide focused experimental studies, and can reduce the need for trial and error, and accidental discoveries. For example, the candidate *de novo* proteins with a possible structure, a specific molecular function (like hydrolysis, or RNA binding), and a propensity to form condensates, can be experimentally probed for these specific properties. Our sequence space analysis can also identify *de novo* proteins that are likely to adopt more conservedprotein-like properties, as a consequence of evolution. Overall, our study not only broadens our understanding of the dynamic nature of protein evolution but also serves as a guidebook for future experimental studies.

## Materials & Methods

### Dataset curation

We used the sequence datasets from our previous study [Middendorf and Eicholt, 2024]. Specifically, we first obtained 6716 orphan protein sequences from the *Drosophila* clade, and their corresponding evolutionary age, from Heames et al. [2020]. From this dataset, we discarded sequences that were annotated with the same FlyBase ID. Next, we extracted the sequences whose emergence origin was annotated as “*denovo*” (intergenic *de novo* protein) or “*denovo-intron*” (intronic *de novo* protein) by Heames et al. [2020], for further analysis. Out of the 2510 proteins sequences thus obtained, 1481 were annotated as “*denovo*,” while 1029 were described as “*denovo-intron*”. Based on their date of emergence, the *de novo* proteins were classified as young (<5 mya), intermediate (5-30 mya), and old (>30 mya) proteins [Heames et al., 2020, Middendorf and Eicholt, 2024]. In our filtered dataset, the three age groups consisted of 2205, 110, and 195 proteins, respectively. We generated 2507 random sequences with the same distributions of amino acid composition and sequence length, as the 2510 *de novo* sequences set, using a technique used in previous studies [Heames et al., 2023, Middendorf and Eicholt, 2024]. We generated a set of conserved protein sequences with the same sequence length distribution as the *de novo* proteins, by randomly sampling protein sequences from the combined proteome of 12 *Drosophila* species. After removing sequences that were duplicated or were redundant with our set of *de novo* proteins, we obtained a set of 2235 unique conserved proteins.

We performed structure predictions using AlphaFold2 [v2.1.1, Jumper et al., 2021] on the High Performance Computing Cluster, PALMA II (University of Muenster). We used the predictions with the highest mean pLDDT for further analysis. We downloaded AlphaFold2 based structure predictions of conserved *Drosophila* proteins from the AlphaFold Protein Structure database [Varadi et al., 2021] for our initial analyses.

### Molecular Dynamics simulations to refine structure predictions

To analyze the stability of the predicted structures of *de novo* proteins, we performed molecular dynamics (MD) simulations using a previously described method [Ferruz et al., 2022], with minor modifications. We only simulated protein structures with i) less than 30% disorder predicted by flDPnn [Hu et al., 2021], and ii) less than 95% of their residues predicted as α-helices by DSSP [Kabsch and Sander, 1983] (1468 unique proteins). We constructed the MD model and performed the simulations using the HTMD python package [Doerr et al., 2016]. The model systems were built to form solvated all-atom cubic boxes. We centered our proteins at the origin of the simulation box coordinates. We used water as the solvent, and added NaCl ions to neutralize the system. We used the AMBER 14SB force field [Maier et al., 2015] for all simulations. We minimized, equilibirated, and simulated each system for 100 ns, using the ACEMD engine [Harvey et al., 2009] with the default settings in triplicates. We evaluated the simulations with the HTMD [Doerr et al., 2016] and MDAnalysis [Michaud-Agrawal et al., 2011] python packages. We calculated the average RMSD value per trajectory for every replicate simulation for a protein, and in turn calculated a single averaged value from three replicates.

### Identifying similar protein structures using Foldseek

We searched the AlphaFold Protein Structure database [Varadi et al., 2021] clustered at 50% sequence identity (AFDB50), for structures similar to the predicted structures of our *de novo*, random, and conserved proteins, using Foldseek [v.8.ef4e960, van Kempen et al., 2023]. We applied the same filtering criteria our query proteins that we used for the MD simulations. For *de novo* proteins, we used the protein structures refined after 100ns of MD simulation. We downloaded pre-computed AFDB50 database via Foldseek’s database module. We searched for similar structures using the “easy-search” module of Foldseek with the default settings. We did not filter the results or queries based on the pLDDT values. We discarded all hits to proteins within the *Drosophila* clade, to exclude hits to orthologous *de novo* proteins.

To identify and annotate potential known protein structural domains in the *de novo* proteins, we searched the protein data bank database [PDB, January 2024; Berman et al., 2000] for structures that were similar to that of *de novo* proteins (MD-refined). We used Foldseek for this analysis with the same settings as we did before for AFDB50. We discarded hits with a TM-score less than 0.5 [Xu and Zhang, 2010]. We retrieved the annotated ECOD domains of the highest scoring hits, from the ECOD database [Release: 20230309, Cheng et al., 2014] if the structural alignment of the *de novo* protein covered at least 80% of the target structure from the PDB. In all cases, we only used the highest scoring hit out of the three MD replicates for further analysis.

### Predicting protein function using DeepFri

To understand the potential function of *de novo*, and random proteins, we predicted their gene ontology (GO) terms using DeepFRI [Gligorijević et al., 2021]. We used their AlphaFold2 predicted 3D-structures as the input and identified hits with a score *≥* 0.5. We summarized the predicted GO terms to a small list of terms using using REVIGO [Supek et al., 2011], and measured semantic similarity using SimRel [Sæbø et al., 2015]. We visually, identified clusters within the semantic space and annotated them with a term that summarizes the GO terms within them.

### Analysis of *de novo* proteins that form biomolecular condensates

We predicted the potential of *de novo* proteins to form biomolecular condensates, using PICNIC [Hadarovich et al., 2023]. Because PICNIC makes predictions based on metrics derived from AlphaFold2 and IUPred2A predictions, we applied further filtering steps of the results in order to obtain a set of high-confidence condensate-forming *de novo* proteins. To this end, we retrieved all the proteins from the CD-CODE database [Rostam et al., 2023], that were experimentally shown from biomolecular condensates *in cellulo* or *in vivo*. This set of 175 proteins served as our positive control. Next, we retrieved sequence features associated with the biological functions of intrinsically disordered regions of proteins [Zarin et al., 2021], using the scripts provided in the idr.mol.feats GitHub repository. We discarded the specific features – *aromatic_spacing*, *omega_aromatic\**, and *kappa\**, and features that count the appearance of specific binding motifs. We normalized all the features that are directly influenced by the sequence length (e.g. amino acid counts), to the sequence length of the corresponding proteins. We subsequently clustered the sequences based on the computed features using hdbscan [McInnes et al., 2017] with a minimal cluster size of 100 the *min_samples* parameter set to a value of 50. We considered a *de novo* protein to be a highconfidence condensate-forming protein, if it shared a cluster with a large fraction of proteins from the CD-CODE database, and had a PICNIC score *≥* 0.5.

### Mapping protein sequences to a numerical space using protein language model

To understand how *de novo* and random protein sequences are located within the protein sequence space relative to conserved proteins, we used the protein language model ESM2 with 650 million parameters (ESM2-650M) [Lin et al., 2023]. Specifically, we used the language model to convert each sequence to a numerical vector with 1280 elements. More specifically, ESM2-650M assigns each amino acid residue in a protein sequence, a 1280-dimensional vector of “embeddings”. For each protein we calculated the multivariate mean of the embedding vectors from every amino acid residue. We calculated the Manhattan distance (or L1 norm) between the numerical sequences of every pair of proteins in our combined dataset of *de novo*, random and conserved proteins. We applied Mann-Whitney test to the pairwise distances to analyse if proteins of one class (*e.g. de novo*) are farther from that of another class (*e.g.* conserved), than with each other. For proteins of one class, we also used the pairwise distances to identify the nearest neighboring protein from the other class. To visualize the location of different proteins in the sequence space, we used UMAP to project and visualize the proteins (numerical sequence) in a two dimensional space [V 0.5.3, McInnes et al., 2018]. We used UMAP with the default settings (n_neigbours = 15, min_distance = 0.1), except for choosing Manhattan distance as the distance metric and optimizing the low dimensional embedding for 5000 instead of 200 epochs.

### Data & statistical analysis

We analyzed most data with Python programming language (v3.9.18) [Van Rossum and Drake, 2009], using the following packages: Pandas (v1.5.3) [Reback et al., 2022], NumPy (v1.26) [Harris et al., 2020], SciPy (v1.11.3) [Virtanen et al., 2020], and BioPython (v1.80) [Cock et al., 2009]. We generated the plots using Matplolib (v3.4.3) [Hunter, 2007]. We performed the Pearson’s *χ*^2^-tests using the “chisquare” from the scipy.stats package. We analyzed protein sequence space with Julia programming language using the packages Distances.jl (v0.10.11) and HypothesisTests.jl (v 0.11.0)

## Acknowledgements

We thank Alun Jones for his advice on statistical tests.

## Supporting information

Supporting information is available on Zenodo 10.5281/zenodo.10557890.

## Code and Data Availability

Datasets are publicly available on Zenodo. All scripts are freely available on GitHub: https://github.com/LasseMiddendorf/SequenceAndFunctionalSpaceOfDrosophilaDeNovoProteins

## Supplementary Material

**Figure S1:**
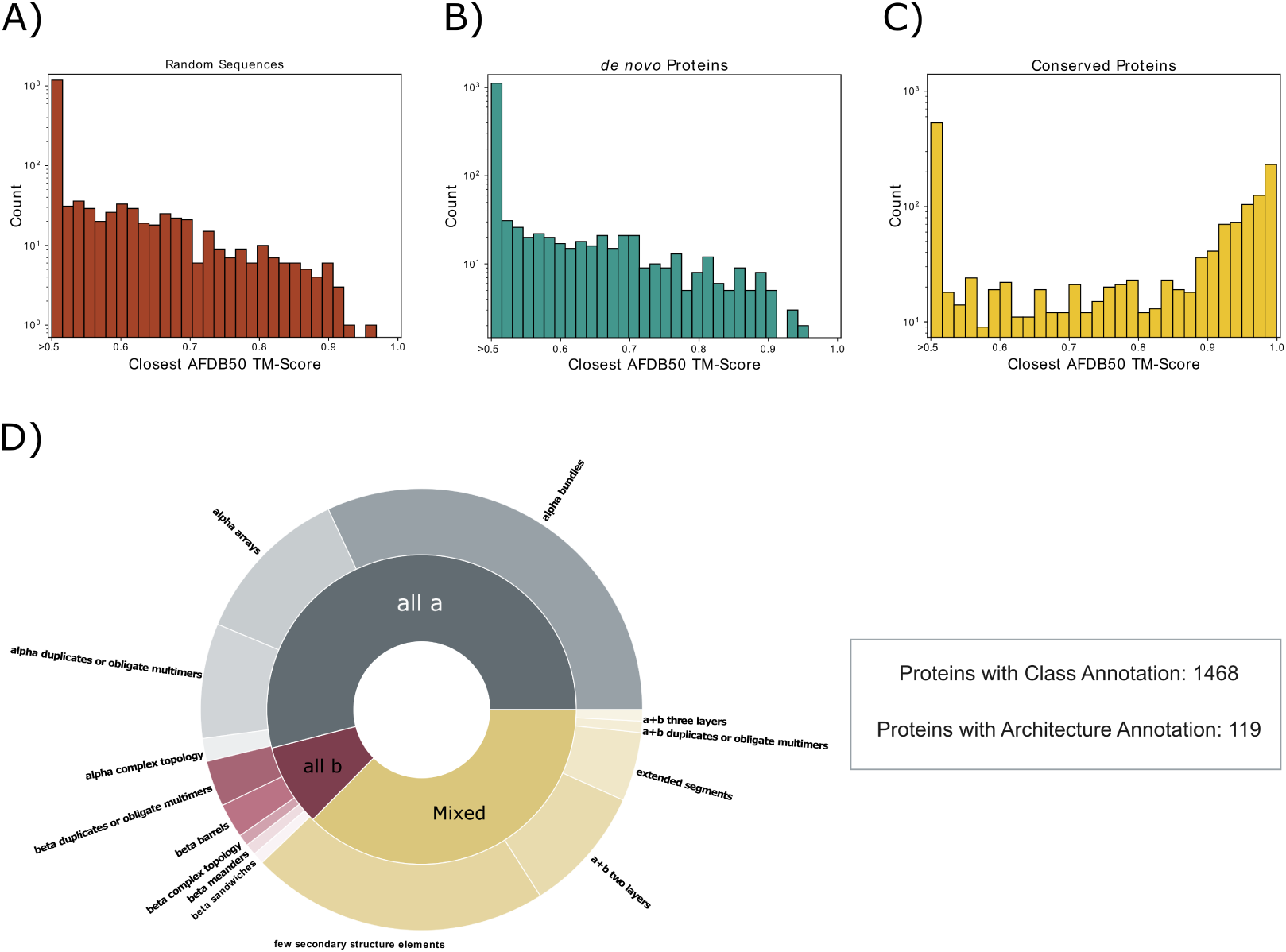
Structural diversity of *de novo* proteins, before MD refinement. The predicted protein structures of randomly generated sequences (**A**), *de novo* protein (**B**), and conserved proteins (**C**) were queried against the AlphaFold database (AFDB50) excluding proteins from *Drosophila*. Only proteins with less than 30% of their residues being predicted to be disordered and less than 95% with a DSSP annotation of being α-helical were considered for the analysis. Shown is the distribution of the highest TM-score found for each protein in the three datasets. (**D**) Overview of the structural classes and ECOD architectures of *de novo* proteins. The protein class (inner circle) was assigned to all *de novo* proteins queried against the AFDB50 based on the DSSP annotations of the predicted protein structures. Proteins containing no residues annotated as α-helices or β-sheets were classified as *all b* or *all a*, respectively. Protein structures containing residues annotated as α-helices and β-sheets were classified as *Mixed*. For the annotation of ECOD architectures in the predicted structures of *de novo* protein, the structures were queried against the PDB and assigned with the ECOD domain of the highest ranking hit if the alignment covered at least 80% of the target structure.

**Figure S2:**
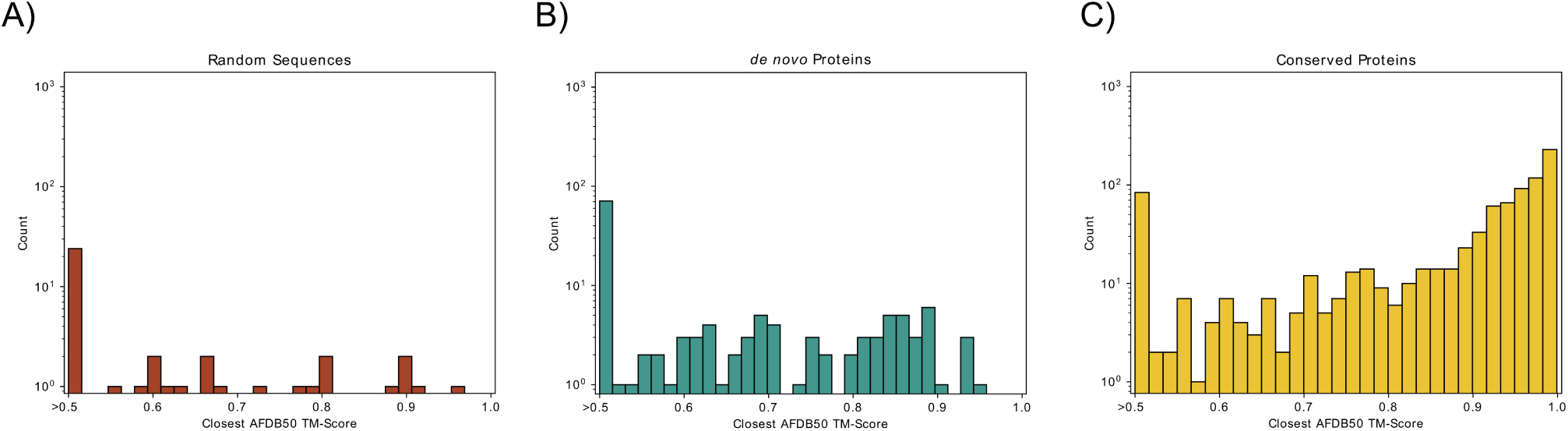
Structural similarity of high-pLDDT protein structures to AlphaFold database. Similar structures in the AlphaFold database for high-pLDDT structure predictions only TM-Score distribution of predicted protein structures of (**A**) random, (**B**) *de novo*, and (**C**) conserved proteins with a pLDDT value >= 70 queried against the AlphaFold database (AFDB50) using Foldseek. The hit with the highest TM-score was chosen for each protein.

**Figure S3:**
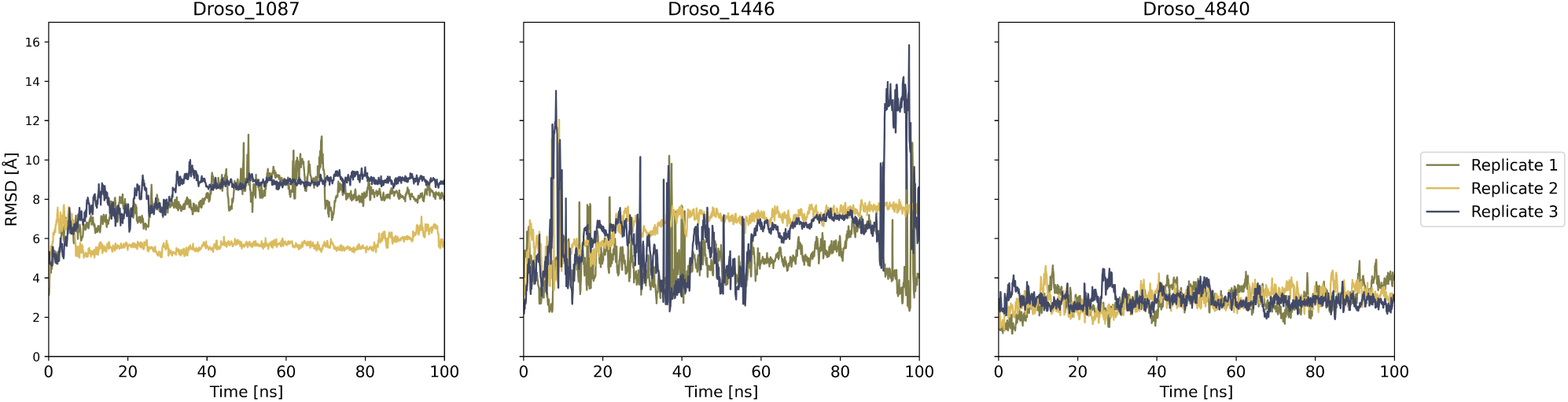
RMSD trajectories of selected *de novo* proteins. Root mean square deviation (RMSD) of Droso_1087, Droso_1446, and Droso_4480 over 100 ns of molecular dynamics simulations. Simulations were performed as triplicates for all proteins.

**Figure S4:**
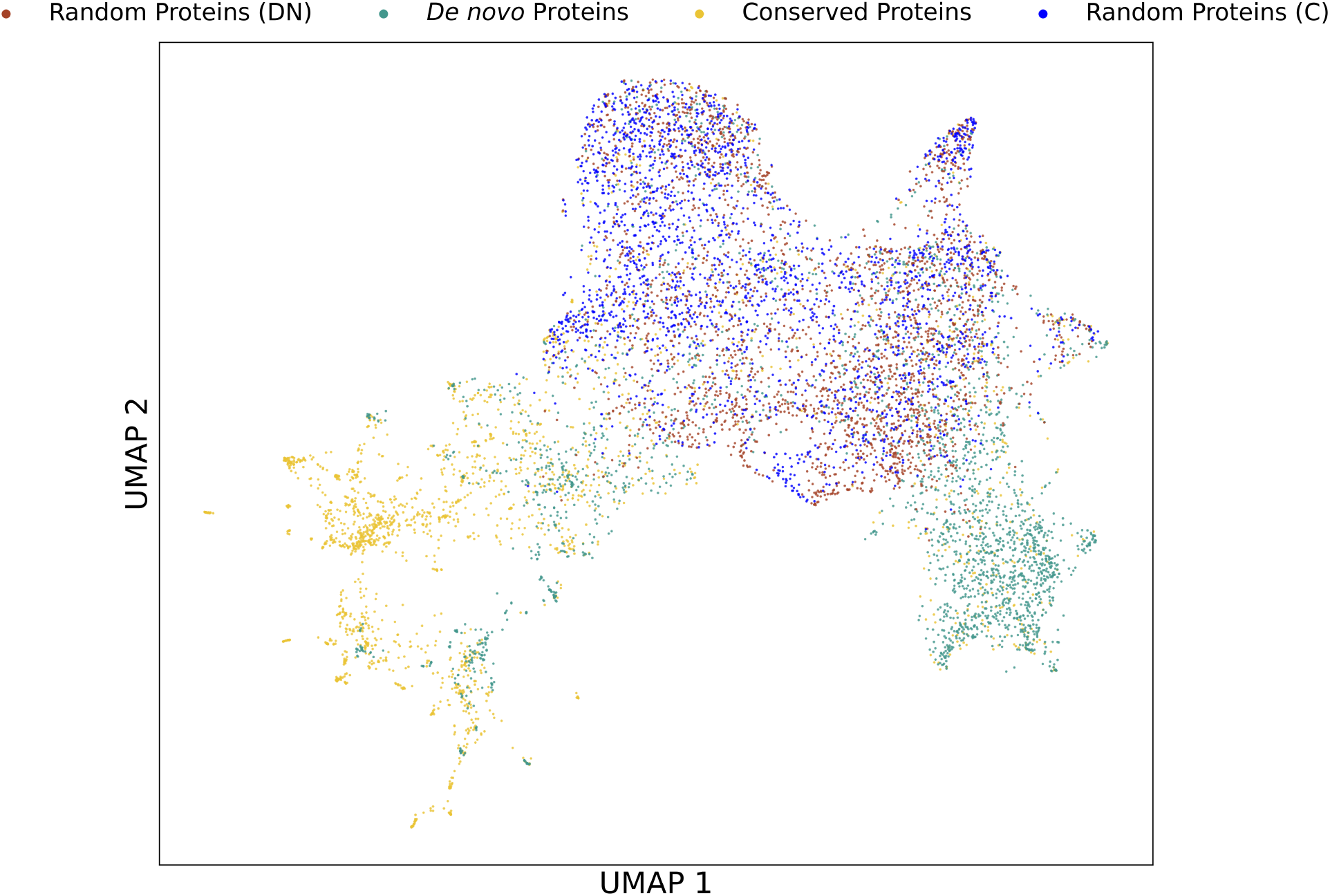
*De novo* proteins are closer to random sequences than conserved proteins. We used the protein language model ESM2-650M to represent the sequences of *de novo*, conserved, and random proteins as numerical vectors. In addition to random proteins that were generated to share the same amino acid distribution as *de novo* proteins (Random Proteins (DN)), we included a set of randomly generated sequences based on the properties of conserved proteins (Random Proteins (C)). We projected the representations into two dimensions using UMAP. The localization in sequence space shows that *de novo* proteins are closer to random proteins than conserved ones, regardless of the origin of the random sequences.

## Notes

### Competing Interest Statement

The authors have declared no competing interest.

